# NAP (davunetide) preferential interaction with dynamic 3-repeat Tau explains differential protection in selected tauopathies

**DOI:** 10.1101/440941

**Authors:** Yanina Ivashko-Pachima, Maya Maor, Illana Gozes

## Abstract

The microtubule (MT) associated protein Tau is instrumental for the regulation of MT assembly and dynamic instability, orchestrating MT-dependent cellular processes. Aberration in Tau post-translational modifications ratio deviation of spliced Tau isoforms 3 or 4 MT binding repeats (3R/4R) have been implicated in neurodegenerative tauopathies. Activity-dependent neuroprotective protein (ADNP) is vital for brain formation and cognitive function. ADNP deficiency in mice causes pathological Tau hyperphosphorylation and aggregation, correlated with impaired cognitive functions. It has been previously shown that the ADNP-derived peptide NAP protects against ADNP deficiency, exhibiting neuroprotection, MT interaction and memory protection. NAP prevents MT degradation by recruitment of Tau and end-binding proteins to MTs and expression of these proteins is required for NAP activity. Clinically, NAP (davunetide, CP201) exhibited efficacy in prodromal Alzheimer’s disease patients (Tau3R/4R tauopathy) but not in progressive supranuclear palsy (increased Tau4R tauopathy). Here, we examined the potential preferential interaction of NAP with 3R vs. 4R Tau, toward personalized treatment of tauopathies. Affinity-chromatography showed that NAP preferentially interacted with Tau3R protein from rat brain extracts and fluorescence recovery after photobleaching assay indicated that NAP induced increased recruitment of human Tau3R to MTs under zinc intoxication, in comparison to Tau4R. Furthermore, we showed that NAP interaction with tubulin (MTs) was inhibited by obstruction of Tau-binding sites on MTs, confirming the requirement of Tau-MT interaction for NAP activity. The preferential interaction of NAP with Tau3R may explain clinical efficacy in mixed vs. Tau4R pathologies, and suggest effectiveness in Tau3R neurodevelopmental disorders.

## Introduction

Microtubules (MTs) are the major component of the neuronal cytoskeleton, and MT stability and organization play a critical regulatory role during axonal transport and synaptic transmission (1). The MT-associated protein Tau is widely expressed in neurons and serves as a primary protein marker for axons (2, 3). Tau promotes MT assembly and regulates MT dynamic instability, which is essential for establishing neuronal polarity, axonal elongation, and neural outgrowth (4). Neurodegenerative disorders with Tau involvement are referred to as tauopathies (5). The Tau protein consists of an N-terminus region projecting outward from the MTs and a C-terminus part directly interacting with the MTs through MT-binding domains (6). Tau3R and 4R (containing either three or four MT-tubulin - binding repeats, respectively) are produced by alternative splicing around exon 10 of the Tau transcript (7). The healthy human brain exhibits a 1/1 ratio of Tau3R/4R and deviation from this ratio are the pathological feature of several tauopathies (8). Phosphorylation of Tau protein controls its binding to MT and is associated with Tau aggregation in neurodegenerative diseases (5,9). In general, Tau3R has been linked to neurodevelopment (7), while Tau4R with aging (10).

We have previously shown that the expression of activity-dependent neuroprotective protein (ADNP), a protein vital for brain formation (11, 12)), is correlated with Tau3R expression (13) and Adnp+/- mice exhibit tauopathy features - significant increase in phosphorylated Tau, prevented by treatment of ADNP-derived peptide NAP (NAPVSIPQ) (14) as well as and tangle-like structures. Our cell culture results have indicated that NAP enhances Tau-MT interaction in the face of zinc intoxication (15) and NAP protective activity requires Tau expression (16). We have further revealed that NAP-Tau association is mediated by direct interaction of NAP and Tau with MT end-binding proteins (EBs) (15, 17).

Clinical trials identified the potential efficacy of NAP (davunetide, CP201) in enhancing short term memory in amnestic mild cognitive impairment patients (18). However, it was not found to be an effective (though, well tolerated) treatment for progressive supranuclear palsy (PSP) patients (19). Because abnormal aggregation of Tau4R is a hallmark of PSP pathophysiology (20), the current study aimed to determine whether NAP had a different activity on either Tau3R or 4R. Our results now showed that NAP preferentially interacted with Tau3R protein from rat brains and induced increased recruitment of human Tau3R to MTs under zinc toxic condition in comparison to Tau4R. Furthermore, we showed that NAP interaction with tubulin was inhibited by paclitaxel obstruction of Tau-binding sites on MTs, confirming the requirement of Tau-MT interaction for NAP activity.

## Results

### Tau from 60-day-old rat brain does not associate with NAP under conditions that Tau from newborn rat brain does

Different tubulin and Tau isotypes are expressed in the course of a rat brain development (10, 21). Newborn-rats predominantly express the Tau3R isoform while adults predominantly express the Tau4R isoform (22). Here, newborn and 60-day-old rat cerebral cortex extracts were analyzed by immunoblotting, and prevailing expression of Tau3R or 4R was observed in the newborn or 60-day-old cortex protein lysate, respectively, as expected (Fig 1A). Then, newborn and 60-day-old rat cortex protein lysates were exposed to NAP affinity chromatography, and eluted proteins were analyzed by immunoblotting with Tau3R and 4R, total-Tau (identifying all Tau isoforms), tubulin Tub2.1 (identifying neuronal-enriched tubulin (23)) and TubpβIII (identifying neuronal-specific tubulin (24)) antibodies. Immunoreactivity for all tested antibodies was detected in the acid eluted fraction from newborn rat brain extract (Fig 1B). However, no significant Tau or tubulin-like bands were observed in the eluted fractions from the mature rat extract under the current experimental conditions (Fig 1B).

**Fig 1.**
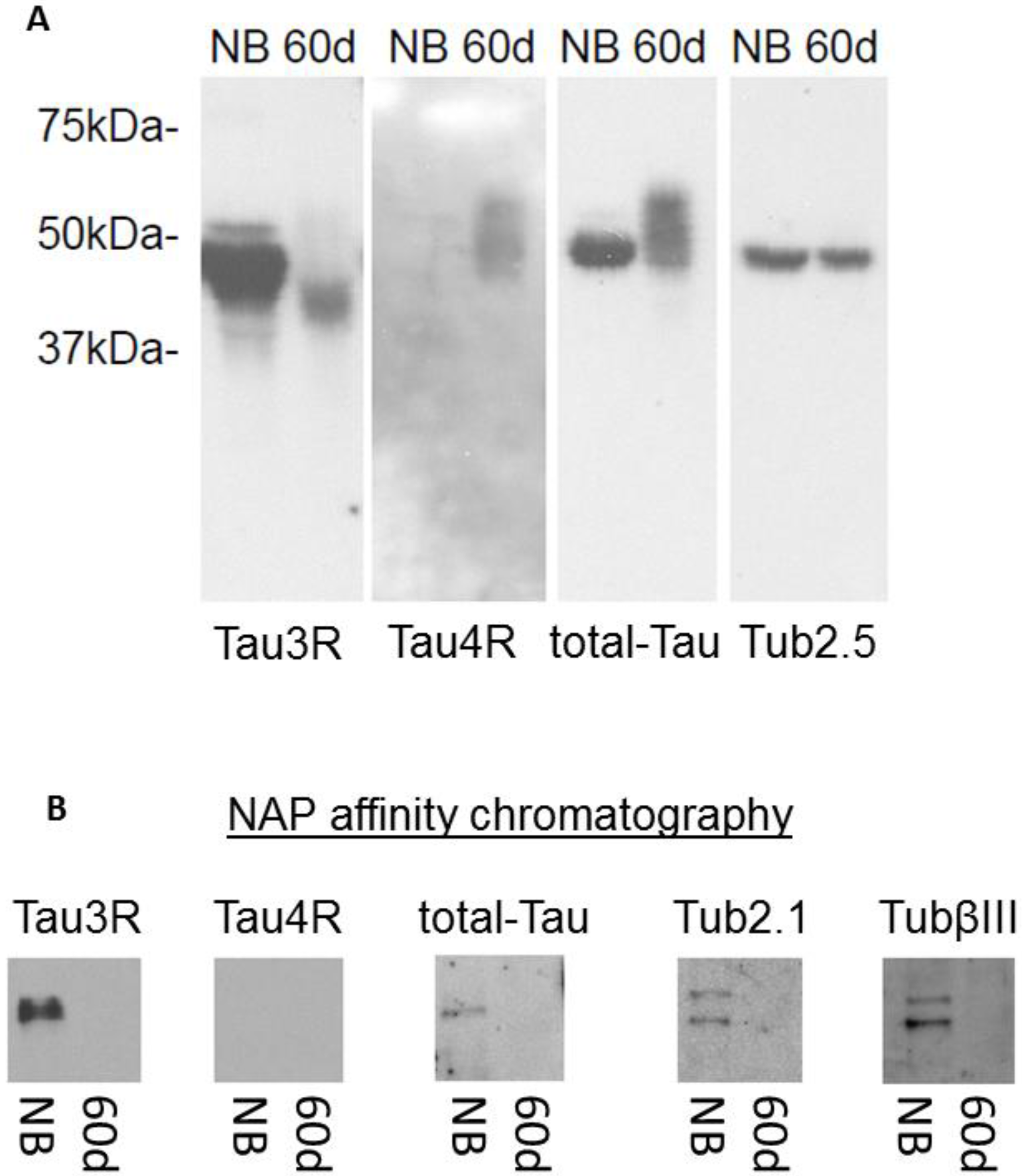
NAP preferentially associates with 3R-tau. **(A)** Western blot analysis of equal amounts of protein extracts from newborn (NB) rat brain cortex and 60-day-old (60d) rat brain cortex. Significantly larger amount of Tau3R was detected in the one-day-old cortical extract compared to the 60-day cortical extract, and Tau4R was recognized in the 60-day cortical extract only. **(B)** Western blot analysis of elution fractions obtained by NAP-affinity chromatography with the same protein extracts of rat brain cortex. Tau3R, total-Tau, and tubulin were identified in the NAP-binding fraction of the newborn rat cortical brain extract, but essentially neither Tau nor tubulin was identified in the elution fraction of the 60-day-old cortical brain extract (three independent experiments).

### NAP induces increased recruitment of human Tau3R to MTs under zinc toxic condition in comparison to Tau4R

In order to test the effect of NAP on the interactions of different Tau isoforms with MTs, fluorescent recovery after photobleaching (FRAP) assay was performed (Fig 2). mCherry-tagged human Tau3R and 4R proteins (S1 Fig.) were over-expressed in differentiated neuroblastoma N1E-115 cells and extracellular zinc (400μM, 1 hour) was used as a MT disruptor, inducer of Tau release from MTs (15). In general, after photobleaching of the region of interest (ROI), the unrecovered portion of initial fluorescence intensity within a bleached area is referred to the immobile fraction of bleached mCherry-Tau proteins because it does not release binding sites on MTs for the entry of un-bleached mCherry-Tau molecules and thus does not allow fluorescence recovery. Therefore, the immobile mCherry-Tau fraction represents MT-bound Tau and reflects the MT-Tau interaction. Here, we observed that treatment with extracellular zinc increased fluorescence recovery 87sec after photobleaching of both mCherry-Tau3R and 4R molecules (Fig 2A). Analysis with one-phase exponential association showed a significant decrease of Tau3R and 4R immobile fractions (Fig 2B and C). NAP added together with zinc decreased fluorescence recovery (Fig 2A) and thus significantly enhanced the immobile fraction of Tau3R and 4R compared to treatment with zinc alone (Fig 2B and C). However, while the Tau4R immobile fraction was restored to untreated control level, while the immobile fraction of Tau3R was further increased in comparison to control values and the difference between Tau3R and 4R immobile fractions was found statistically significant (Fig 2C). Thus, NAP treatment produces a more potent impact on 3R-, rather than on 4R-Tau association with MTs.

**Fig 2.**
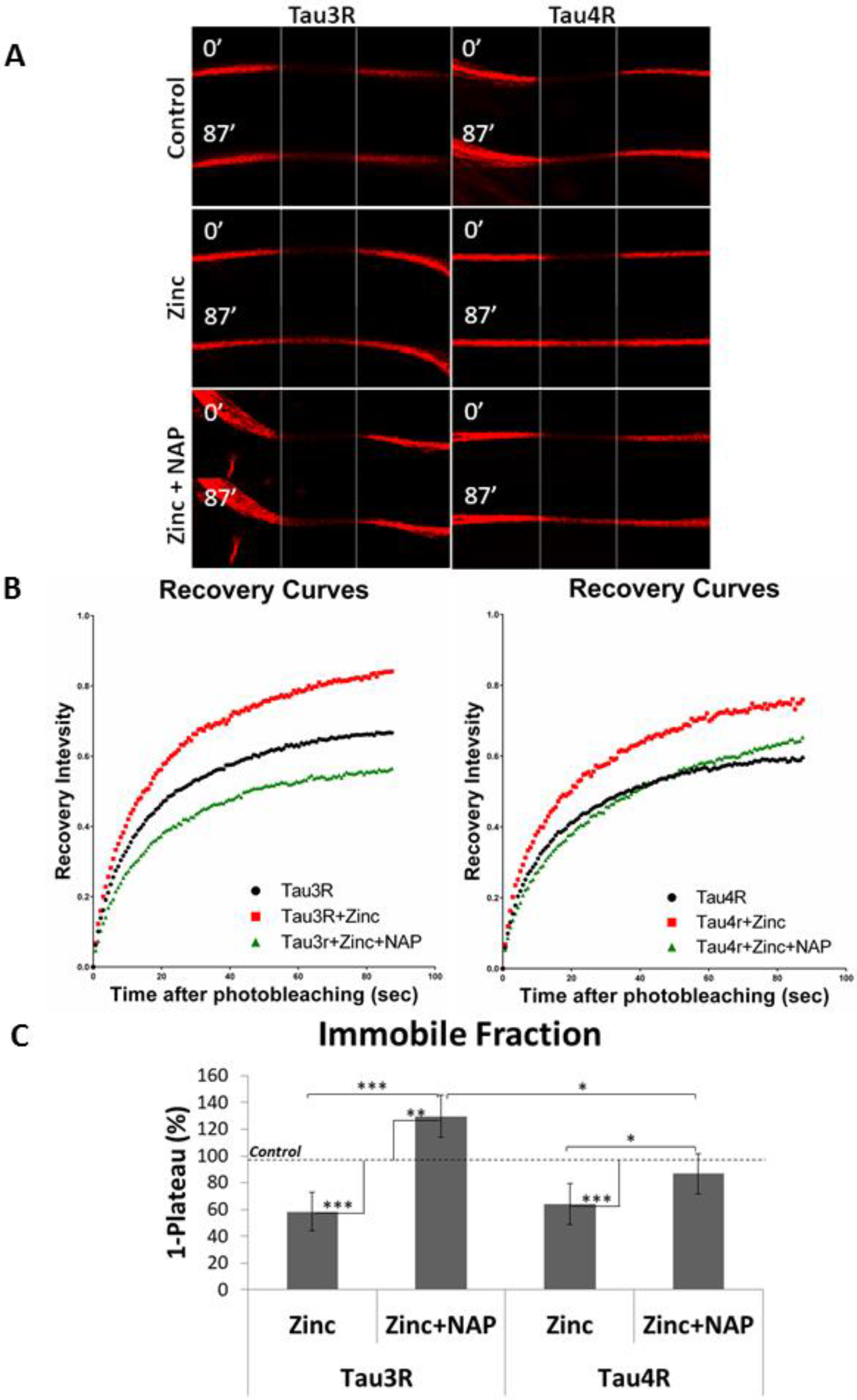
NAP induces increased recruitment of human Tau3R to MTs under zinc toxic condition in comparison to Tau4R. **(A)** Representative images of photo-bleaching and fluorescence recovery of mCherry-tagged Tau3R and 4R in differentiated N1E-115 cells treated with extracellular zinc (400μM, 2hrs) with or without NAP treatment (10^−12^M, 2hrs). N1E-115 cells expressing m-Cherry-Tau3R/4R without any treatment represented the control. **(B)** FRAP recovery curves of normalized data (see “Materials and Methods”). **(C)** The graph represents percentages (±SEM) of the fitted data (from three independent experiments) of immobile fractions relative to control – 100%. Normalized FRAP data were fitted with one-exponential functions (GraphPad Prism 6), and statistical analysis was performed by Two Way ANOVA (SigmaPlot 11). Statistical significance is presented by *P<0.05, **P<0.01, *** P<0.001. Tau3R: Control n=58, zinc n=85, zinc + NAP n=58; Tau4R: Control n=56, zinc n=47, zinc + NAP n=60.

### Tau interaction with MTs is required for NAP activity

Further, we aimed to test requirement of Tau-MT association for NAP interaction with tubulin/MTs. Because paclitaxel obstructs Tau-binding sites on tubulin (25) we incubated NAP-affinity columns with newborn cerebral cortical extracts in the presence of paclitaxel dissolved in DMSO or in the presence of DMSO alone (used as a control). Paclitaxel markedly decreased tubulin immunoreactivity in the acid-eluted fractions compared to the control (Fig 3A and B). Immunoreactivity of Tau3R appeared in both fractions – with and without paclitaxel exposure (Fig 3C). As tubulin was washed away in the presence of paclitaxel, whereas Tau3R remained bound to the NAP column, we suggest that NAP interaction with tubulin required mediation of Tau (Fig 3D). To ascertain the specificity of NAP binding, an affinity control column with eight-amino-acids inactive peptide (VLGGGSALL) was prepared, as well. The peptide VLGGGSALL has previously shown no MT-related neuroprotective activity (26). Affinity chromatography with the control peptide showed Tau3R presence in the loaded material, column flow-through, and column wash, but did not detect Tau3R in the acid elution fractions of both columns, neither in the absence no in the presence of paclitaxel (S2 Fig). Also, tubulin was associated with VLGGGSALL regardless of paclitaxel presence, demonstrating nonspecific interaction (S2 Fig).

Then, we assessed the protective activity of NAP against increased concentrations of paclitaxel. For this purpose, differentiated N1E-115 cells were exposed to paclitaxel (5, 6 and 7 μM) with or without NAP (10^−15^, 10^−12^, 10^−9^M) for 4hrs. Cell viability, measured by mitochondrial activity, was significantly reduced following the 4hr-incubation period with paclitaxel. However, co-treatment with NAP (10^−12^ and 10^−9^M, but not 10^−15^M) protected against the lowest tested concentration of paclitaxel - 5μM (Fig 4), but not against increased concentrations of paclitaxel – 6 and 7μM. These results suggest a requirement of direct Tau-MT interaction for NAP activity, confirming our previously published data that showed requirement of Tau expression for NAP protective capabilities (16).

**Fig 3.**
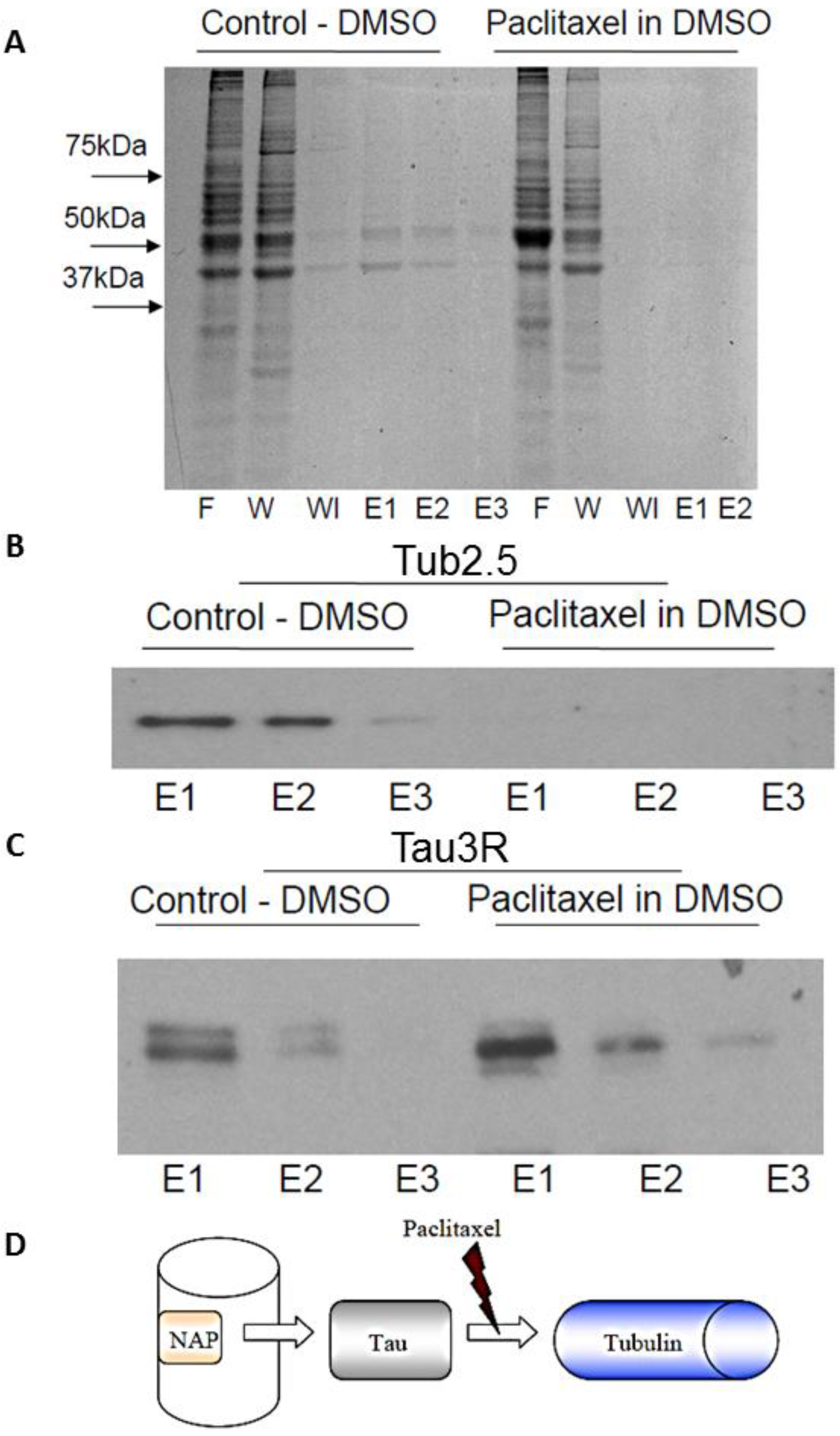
NAP interaction with tubulin, but not with Tau, is inhibited by paclitaxel. **(A)** Bio-SafeTM Coomassie protein staining of the different fractions (F – flow-through, W - first wash, Wl - last wash, E1/2/3 – elution fractions by order) obtained from NAP-affinity column loaded with protein extracts of newborn rat cerebral cortex with 4 mg paclitaxel, dissolved in 80μl DMSO, or equal volume of DMSO, alone. Almost no tubulin is evident in the elution fractions in the paclitaxel column, in contrast to the control one. (**B, C**) Western analysis of elution fractions obtained similarly as in panel (A). **(B)** Tubulin is not detected in the elution fractions of the pre-incubated paclitaxel column in comparison to the control column. **(C)** Tau3R is observed in the elution fractions of both the pre-incubated paclitaxel column and the control column. **(D)** Graphic depiction of the hypothesis that NAP, bound to an affinity column, interacts with tubulin throughout mediation of Tau.

**Fig 4.**
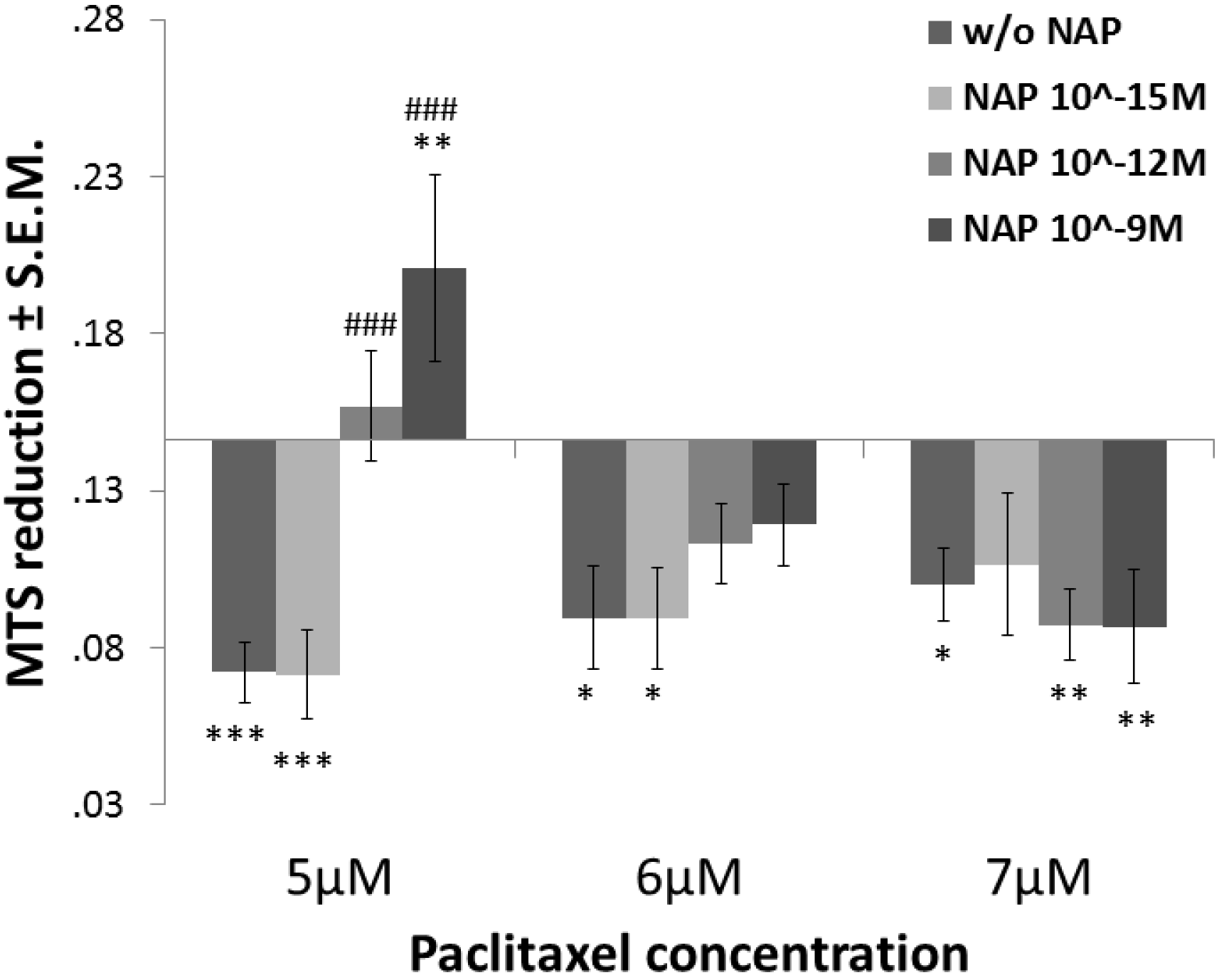
NAP protective activity is inhibited by paclitaxel. Cell viability test performed by the MTS assay (measures mitochondrial activity, see “Materials and Methods”). Differentiated N1E-115 cells are exposed to different concentrations of NAP (10^−15^M, 10^−12^M, 10^−9^M) and paclitaxel (5, 6 and 7 μM) for 2 hrs. Average of mitochondrial activity results (MTS reduction) are displayed in relation to the control (non-treated cell) value – 0.1411±0.02989. Statistical analysis of the data was performed using one-way ANOVA with Tukey post hoc test, n=5. Statistical significance is presented relative to control as *P<0.05, **P<0.01, *** P<0.001; to “w/o NAP” (cells treated with paclitaxel, alone) as #P<0.05, ##P<0.01, ###P<0.001.

## Discussion

As dynamic tracks for motor proteins, MTs are involved in axonal transport and synaptic transmission. We have previously shown that NAP provides neuroprotection (26, 27) and neurotrophic activities (28) through interaction with MTs (15), rescues impaired axonal transport (29–31), regulates dendritic spines (29, 32, 33) and enhances memory (33). NAP also protects against the accumulation of pathologically modified, hyperphosphorylated Tau (14, 34–36). We have previously suggested the mechanism of NAP protective activity on MT-mediated cellular processes through the involvement of Tau and MT end-binding proteins (EBs) (16, 29). Specifically, NAP contains an ADNP association site (SIP) a signature motif for direct interaction with the EB1 and the EB3 proteins (29), which in turn bind to MTs (37) and Tau (17). Furthermore, Tau has been identified as a regulator of EB’s action, and localization on MTs in developing neuronal cells (17) and NAP increases Tau-EB1/3 association (16). Tau is essential for the establishment of MT dynamic instability and axonal transport, while EB1 is more prevalent in neuronal axons (38) and EB3 in dendritic spines (39). Formerly, it has been shown that expression of Tau and EB1/3 proteins are required for NAP-dependent neuronal survival (16, 29). Here, we added details to the understanding of the molecular mechanism underlying MT-related activity of NAP. Our current experiments showed that NAP preferentially interacted with rodent Tau3R and induced enhanced recruitment of human Tau3R to MTs under zinc toxic condition in comparison to Tau4R. Furthermore, we demonstrated that paclitaxel-disturbed Tau-tubulin interaction prevented NAP association with tubulin/MTs and inhibited NAP protective activity, suggesting the requirement of not only the expression of Tau, but also Tau-tubulin direct association for sufficient action of NAP.

The only difference between the two Tau isoforms (3R and 4R) is the presence of the exon 10 coding sequence comprising an extra MT-binding repeat in Tau4R (Fig 5A, red sequence) which is excluded during alternative splicing in Tau3R (40). Elm prediction analysis (41) of the whole Tau sequence identified cyclin A-docking motif within the translated sequence of exon 10 (Fig 5A, S1 Table). Whereas cyclin-dependent kinase (Cdk) 5 is activated by non-cyclin proteins, Cdk1/2 requires direct association with cyclin A imposing an active conformation on the kinase (42). Furthermore, six docking/phosphorylation sites of the cyclin-dependent kinase subunit 1 (Cks1) were identified by Elm prediction on Tau (Fig 5A, S1 Table). Cks1 associated with Cdk-cyclin complex increases the specificity and efficiency of Cdk substrate phosphorylation (43). It has been reported that Cdk2 and Cdk5 provide different Tau phosphorylation profiles (44). It was further reported that region-specific Tau phosphorylation might attenuate Tau-EB association (45). Because NAP interacts with Tau through EB proteins, and Tau3R and 4R may present differences in the phosphorylation profiles, we speculate that observed attenuation of NAP-Tau4R interaction occurs due to some phosphate incorporation on Tau and ensuing decrease of EB protein association (Fig 5B). In this respect, we have previously reported that NAP reduces Tau phosphorylation at Ser262 (30), Ser202/Thr205, and Thr231 (46) residues, but does not exhibit a significant impact on Tau phosphorylation level at Thr181 (19). Intriguingly, Thr181 is on one of the predicted Tau binding/phosphorylation motifs of Cks1 modulating the activity of Cdk (Fig 5A, S1 Table).

As opposed to rodents, Tau3R is abundant alongside with Tau4R in the human adult brain. The ratio of Tau 3R:4R proteins is important, and changes in the ratio are observed in tauopathies (47). NAP does not interact with MT proteins from various cancer cell lines or fibroblasts (27), which may not express any MT-associated proteins with properties that are similar to Tau3R, and NAP does not affect cell division (48). Furthermore, NAP does not protect cells from fibroblast origin unless those cells are transfected with Tau3R-expressing plasmid (16). However, NAP protects MT organization in mature neurons and glia (26, 27, 49). We have previously shown that by promoting the interactions of Tau and EBs with MTs, NAP protects MTs against degradation and concurrently enhances MT dynamics (16). *In vivo*, chronic NAP treatment reduces excess Tau accumulation and pathological hyperphosphorylation (14, 35, 46). Our studies explain, in part, the efficacy of NAP (davuntide, CP201) in enhancing cognitive functions in mild cognitive impairment patients (18), while showing no efficacy (although high safety) in the 4R tauopathy progressive supranuclear palsy (PSP) (19). Since tauopathy underlies a verity of neurodegenerative conditions, our current findings may pave the path for treatment by peptide drugs that have an impact on tubulin-Tau interaction and specific neurofibrillary tangle populations (50). As neurodevelopment has been linked to Tau3R (7, 10), our results pave the path to the development of NAP (davunetide, CP201) for ADNP deficiencies associated with neurodevelopment, for example, the autism-like ADNP syndrome, resulting from de novo truncating mutations in ADNP (32).

**Fig 5.**
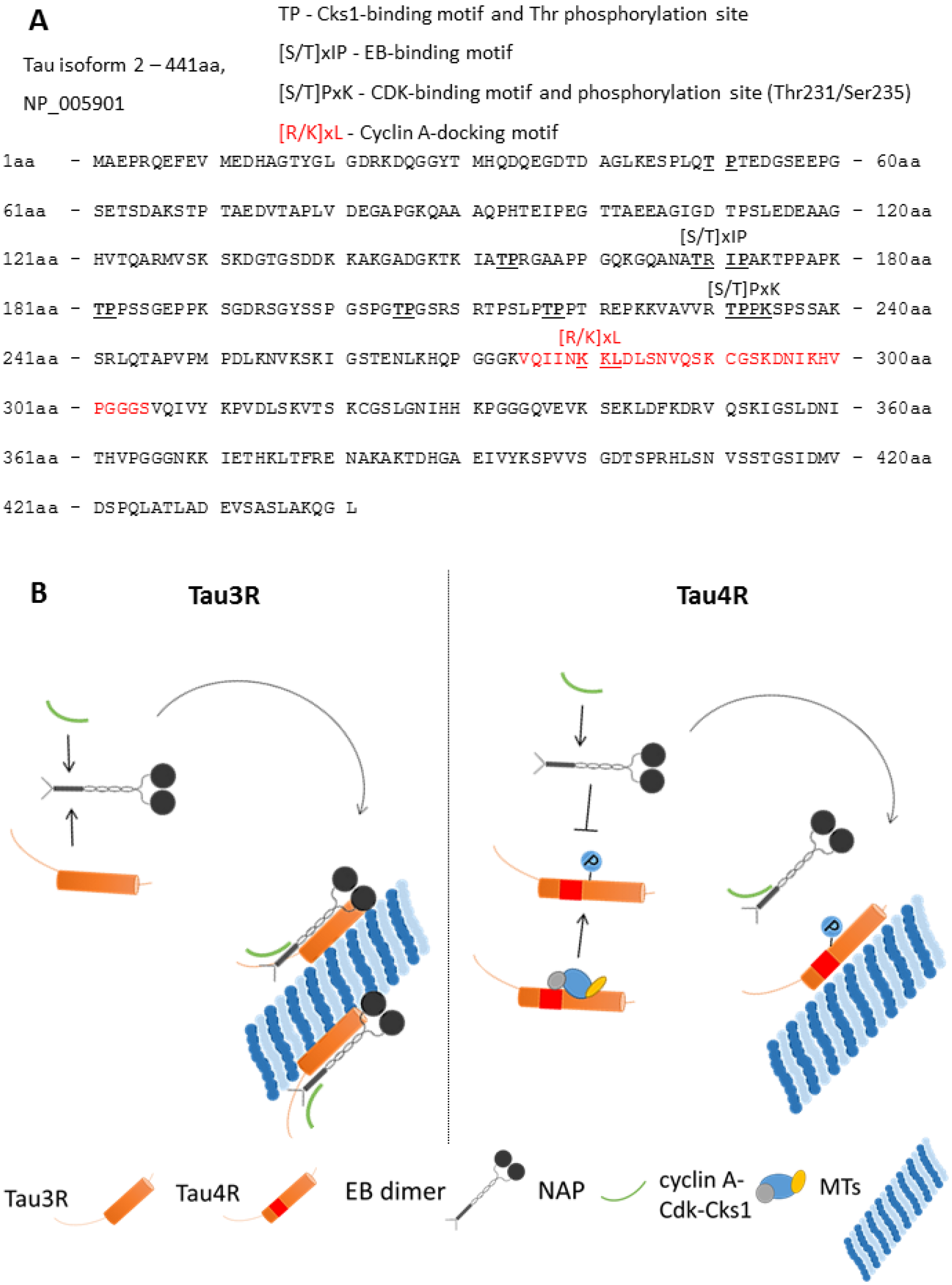
Suggested explanation for the preference of NAP binding to Tau3R over Tau4R. **(A)** Amino-acid sequence of the Tau isoform 2 (NP_005901, 441aa) (Tau4R). The translated protein sequence of exon 10, which is spliced in Tau3R, is marked by red. Functional motifs, predicted by Elm analysis (41) (S1 Table) are indicated. **(B)** Graphic depiction of speculated hypothesis suggesting the preference of NAP to interact with Tau3R based on the findings of Elm prediction analysis. The second MT-binding repeat of Tau4R (spliced in Tau3R) includes cyclin A-docking motif and thus may enable Tau binding to cyclin A, essential for the activation of Cdk1/2 (42). Cks1 may also associate with Cdk and cyclin A to form more efficient cyclin A-Cdk-Csk1 phosphorylation complex. Tau4R phosphorylation by Cdk1/2 differs from Cdk5 (a conventional Tau kinase) (44) and may disturb/attenuate Tau-EB interaction, which has been previously indicated as a crucial for NAP interaction with MTs (16, 29). The EB dimer structure was constructed according to a published review (37).

## Materials and Methods

### Brain extract preparation

Protein lysate was prepared from either one- or sixty-day-old rat cerebral cortex in a lysis buffer containing: 150 mM NaCl, 1 mM EDTA, 50 mM Tris-HCl, pH 4.5 (or 7.5 as indicated), 0.1% Triton X-100, 1% Nonidet P-40, and a protease inhibitor cocktail (Roche Diagnostics, Mannheim, Germany). DNA was fragmented by sonication. Cell debris was discarded following 20 minutes of centrifugation at 30,000g at 4°C, as described previously (26).

### NAP affinity chromatography

Affinity columns contained extended NAP (**CKKKGG**NAPVSIPQ, the linker peptide is in bold). Peptides were purchased from Genemed Synthesis, Inc., San Antonio, TX, USA or synthesized as before (51). Columns and peptide binding were prepared using Sulfolink coupling gel (Pierce, Rockford, IL, USA) according to the manufacturer’s instructions as before (26). 2 ml Sulfolink coupling gel was loaded onto Poly-Prep Chromatography Columns (Bio-Rad, Hercules, CA, USA) and coupled with 2mg/ml peptide. Coupling was ascertained by free peptide measurements.

In order to determine the binding specificity of Tau and tubulin to NAP, multiple experiments were performed under stringent comparable experimental conditions as follows:

1] Proteins from either newborn or sixty-day-old rat cerebral cortical extracts (2mg protein/ml, total volume 2 ml) were loaded onto the NAP-affinity columns at pH 7.5 and incubated for 16 hours at 4°C, columns were washed with phosphate buffered saline (PBS, 20-25 ml) until all unbound protein had eluted as confirmed by the Bradford protein assay (Bradford, BioRad, Hercules, CA, USA). The bound protein was then eluted with 0.1 M glycine pH 2.6.
2] Proteins from one-day-old rat (expressing Tau3R, solely) cerebral cortical extracts were loaded onto the NAP column with 4 mg paclitaxel (Haorui Pharma-Chem Inc., New-Jersey, USA) dissolved in 80μl dimethyl sulfoxide (DMSO, Sigma, Rehovot, Israel). A control NAP column was treated similarly with 80μl DMSO but without paclitaxel. The columns were incubated with brain extract, washed as described above, and eluted with glycine 0.1 M pH 2.6.
3] The control column contained an inactive peptide **CKKKGG**VLGGGSALL (the linker peptide is in bold) described in supplemental materials and methods (S1 File and S2 Fig).

### SDS-PAGE and western blot analysis

The flow-through, wash fractions and elution fractions were separated by 10% or 12% SDS polyacrylamide gel electrophoresis (SDS-PAGE) followed by protein staining using Bio-SafeTM Coomassie (Bio-Rad, Hercules, CA, USA) according to manufacturer’s instructions or transferred to nitrocellulose membranes (Schleicher and Schull, Dassel, Germany) for western blot analysis. In comparative experiments, run in parallel, the same amounts were loaded on the gels, for each parallel fraction. Non-specific sites on the nitrocellulose membranes used for western analysis were blocked in a blocking solution (10 mM Tris pH 8, 150 mM NaCl, and 0.05% Tween 20 [TBST]) supplemented with 5% non-fat dried milk (1 hour, at room temperature). The Protein complexes were visualized by SuperSignal West Pico Chemiluminescent Substrate (Pierce, Rockford, IL, USA).and exposed on Fuji Film Medical X-ray film (Fuji Corporation, Tokyo, Japan).

### Antibodies

Total-Tau - mouse monoclonal antibody Tau5 (antibody recognizing all Tau forms) was obtained from MBL International Corporation (Woburn, MA, USA). Tau3R and Tau4R - mouse monoclonal anti-Tau RD3 (3-repeat isoform) and mouse monoclonal anti-Tau RD4 (4-repeat isoform) were obtained from Millipore Corporation (Billerica, MA, USA). Tub2.1 and Tub2.5 - mouse monoclonal tubulin antibodies maintained and kindly provided by Professor Colin Barnstable, and were used as before (23). TubβIII - mouse monoclonal antibody β tubulin isotype III was obtained from Sigma-Aldrich (St. Louis, MO, USA). Secondary antibodies were goat anti-mouse-horseradish peroxidase - HRP (Jackson ImmunoResearch, West Grove, PA, USA). All antibodies were used at the dilution of 1:1000, except when otherwise indicated.

### Plasmid construction

DNA inserts carrying Tau3R and 4R were obtained from human Tau3R and 4R cDNA containing plasmids (a kind gift of Professor M. Goedert, MRC Laboratory of Molecular Biology, Cambridge, UK) and then cloned into the backbone of the newly constructed pmCherry-C1 plasmid. For more details, see supplemental materials and methods (S1 File and S1 Fig).

### Cell culture and treatments

Mouse neuroblastoma N1E-115 cells (ATCC, Bethesda, MD; passage numbers from 10 to 13) were maintained in Dulbecco’s modified Eagle’s medium (DMEM), 10% fetal bovine serum (FBS), 2 mM glutamine and 100 U/ml penicillin, 100 mg/ml streptomycin (Biological Industries, Beit Haemek, Israel). The cells were incubated in 95% air/5% CO_2_ in a humidified incubator at 37°C. N1E-115 cells were plated on 35mm dishes (81156, 60 μ-Dish, Ibidi, Martinsried, Germany) at a concentration of 25*10^4^ cells/dish and then were differentiated with reduced FBS (2%) and DMSO (1.25%) containing medium during five days before transfection and seven days before the experiment. On the day of experiment differentiated N1E-115 cells were treated for 1 hrs with zinc chloride (ZnCl_2_; final concentration, 400 μM, Sigma, Rehovot, Israel) with or without NAP (10^−12^M).

### Transfection of over-expression plasmids and Fluorescence recovery after photobleaching (FRAP)

5-day differentiated N1E-115 cells were transfected with a 1μg pm Cherry-C1-Tau3R/4R plasmid. 48 hrs after transfection, cultured N1E-115 cells were incubated at 37 °C with a 5% CO_2_/95% air mixture in a thermostatic chamber placed on the stage of a Leica TCS SP5 confocal microscope [objective 100x (PL Apo) oil immersion, NA 1.4]. An ROI (region of interest) for photo-bleaching was drawn in the proximal cell branches. mCherry-Tau3R/4R was bleached with a 587nm argon laser, and fluorescence recovery was at 610-650nm. Immediately after bleaching, 80 images were collected every 0.74s. Fluorescence signals were quantified with ImageJ (NIH), obtained data were normalized with easyFRAP43, and FRAP recovery curves were fitted by a one-phase exponential association function using GraphPad Prism 6 (GraphPad Software, Inc., La Jolla, CA). Samples with R2<0.9 were excluded.

### Cell viability assay

7-day differentiated N1E-115 cells were treated with different concentrations of NAP (10^−15^M, 10^−12^M and 10^−9^M) and paclitaxel (5, 6 and 7μM diluted in DMSO) for 2 hours. Treatments with 5, 6 and 7μM of DMSO alone were used as controls. Cell viability was measured using the MTS assay (CellTiter 96 AQueous Non-Radioactive Cell Proliferation Assay; Promega, Madison, WI, USA), which was performed according to the manufacturer’s instructions and read in an ELISA plate reader at 490nm.

### Statistical Analysis

Data are presented as the mean ± SEM from 3 independent experiments. Statistical analysis of the data was performed by using one-way ANOVA test (followed by the Tukey post hoc test) by the IBM SPSS Statistics software version 23. * P<0.05, ** P<0.01, *** P<0.001.

## Acknowledgements

This work is in partial fulfillment of the PhD thesis requirements of Msc. Yanina Ivashko-Pachima. Work was supported in part by the following grants, ISF 1424/14, ERA-NET neuron AUTISYN, BSF-NSF 2016746, AMN Foundation as well as Drs. Ronith and Armand Stemmer, Mr Arthur Gerbi (French Friends of Tel Aviv University), and Spanish Friends of Tel Aviv University. Professor Illana Gozes is the incumbent of the Lily and Avraham Gildor Chair for the Investigation of Growth Factors, and the Director of the Dr. Diana and Zelman Elton (Elbaum) Laboratory for Molecular Neuroendocrinology at Tel Aviv University. Professor Gozes also serves as the Chief Scientific Officer of Coronis Neurosciences, developing CP201 for the ADNP syndrome.

## Supporting Information

**S1 Fig. Plasmid maps**. pmCherry-C1 vector **(A)**, human Tau3R **(B)** and 4R **(C)** expressing plasmids based on pmCherry-C1 vector. The plasmid maps were constructed with Benchling platform (www.benchling.com).

**S2 Fig. VLGGGSALL does not interact with Tau3R.** The gel lanes contain the protein loaded (load) flow-through (FT) PBS wash, pH7.5 (W1-30) and acid elutes (E1-E4) from the columns linked to eight-amino-acid inactive peptide VLGGGCALL P (has previously shown no microtubule-related neuroprotective activity (26)) that were incubated with brain extracts and DMSO in the absence and presence of paclitaxel. Western blotting analysis with anti Tau RD3 detected Tau3R presence in the loaded material, column flow-through and column wash, but did not detect Tau3R in the acid elution fractions of both the columns. In contrast, tubulin antibodies - Tub2.5 identified tubulin-like bands also in the elution fractions with no apparent influence of paclitaxel treatment.

**S1 Table. Elm prediction analysis of Tau (NP_005901) exon 10 translation sequence.** Elm analysis (41) predicted functional motifs of the translation sequence of spliced exon 10 (VQIINKKLDLSNVQSKCGSKDNIKHVPGGGS) of Tau isoform 2 (NP_005901). DOC_CYCLIN_RxL_1 motif appeared only once in full Tau sequence.

